# Expression of transcription factors KLF2 and KLF4 is induced by the mammalian Golgi stress response

**DOI:** 10.1101/2023.05.16.541051

**Authors:** Kanae Sasaki, Miyu Sakamoto, Iona Miyake, Reishi Tanaka, Ryuya Tanaka, Azusa Tanaka, Misaki Terami, Ryota Komori, Mai Taniguchi, Sadao Wakabayashi, Hajime Tajima Sakurai, Hiderou Yoshida

## Abstract

The Golgi stress response is a homeostatic mechanism that augments Golgi function when Golgi function becomes insufficient (Golgi stress). Glycosylation of the core proteins of proteoglycans is one of the important functions of the Golgi. If the production of core proteins is increased and the amount of glycosylation enzymes for proteoglycans becomes insufficient (PG-type Golgi stress), the proteoglycan pathway of the Golgi stress response is activated, resulting in the transcriptional induction of glycosylation enzymes, including NDST2, HS6ST1 and GLCE. The transcriptional induction of these glycosylation enzymes is regulated by the enhancer element, PGSE-A; however, transcription factors that induce transcription from PGSE-A have not yet been identified. We herein proposed KLF2 and KLF4 as candidate transcription factors for transcriptional induction from PGSE-A, and revealed that their expression was up-regulated in response to PG-type Golgi stress. These results suggest that KLF2 and KLF4 are important regulators of the proteoglycan pathways of the mammalian Golgi stress response.

## Introduction

Eukaryotic cells contain various organelles, such as the endoplasmic reticulum (ER) and Golgi apparatus. The capacity of each organelle is separately regulated in accordance with cellular demands by the mechanism of organelle autoregulation (Sasaki and Yoshida, 2019). Organelle autoregulation is an indispensable mechanism for eukaryotic cells to function autonomously and is one of the fundamental issues in cell biology (Sasaki and Yoshida, 2015). The ER stress response (also called the unfolded protein response (UPR)) is the organelle autoregulation of the ER, and consists of the following three response pathways: the ATF6, IRE1, and PERK pathways (Costa-Mattioli and Walter, 2020; Marciniak *et al*., 2022; Ninagawa *et al*., 2021; Yong *et al*., 2021; Yoshida, 2007). Each response pathway is regulated by a distinct set of regulators including a sensor, transcription factor, and enhancer element. In the case of the IRE1 pathway, the sensor IRE1 detects ER stress (an insufficiency in ER function) and increases the expression of the transcription factor XBP1, which, in turn, induces the transcription of ER-related proteins by binding to an enhancer element, called UPRE, thereby promoting ER function (Calfon *et al*., 2002; Shen *et al*., 2001; Yamamoto *et al*., 2007; Yamamoto *et al*., 2008; Yamamoto *et al*., 2004; Yoshida *et al*., 2001; Yoshida *et al*., 2009).

One of the functions of the Golgi apparatus is post-translational modifications, such as glycosylation. The autoregulatory mechanism of the Golgi apparatus is called the Golgi stress response (Sasaki and Yoshida, 2019), which has several response pathways, including the TFE3 pathway (Oku *et al*., 2011; Taniguchi *et al*., 2015), HSP47 pathway (Miyata *et al*., 2013), CREB3 pathway (Oh-Hashi *et al*., 2021; Reiling *et al*., 2013), proteoglycan pathway (Sasaki *et al*., 2019), mucin pathway (Jamaludin *et al*., 2019), cholesterol pathway (Kimura *et al*., 2019), GOMED pathway (Noguchi and Shimizu, 2021), GARD pathway (Eisenberg-Lerner *et al*., 2020), ATF4-CSE pathway (Sbodio *et al*., 2018), Golgi-specific transcriptional stress response pathway (Serebrenik *et al*., 2018), and calcium pathway (Smaardijk *et al*., 2018). These response pathways have been suggested to separately augment the capacity of each functional zone in the Golgi apparatus (Sasaki and Yoshida, 2019). The mucin pathway is activated and induces the transcription of glycosylation enzymes for mucins, such as GALNT5, GALNT8, and GALNT18, through the enhancer element mucin-type Golgi stress response element (MGSE) when the synthesis of mucin core proteins is increased and the capacity of glycosylation enzymes for mucins becomes insufficient (mucin-type Golgi stress). In contrast, the proteoglycan pathway induces the transcription of glycosylation enzymes for proteoglycans, including NDST2, HS6ST1, GLCE, B3GAT3, CSGALNACT2, EXT2, HS3ST1, and CHST7, when the production of proteoglycan core proteins is up-regulated and overwhelms the capacity of the respective enzymes (PG-type Golgi stress) (Sasaki et al., 2019). An analysis of the promoters of the target genes of the proteoglycan pathway revealed two enhancer elements responsible for their transcriptional induction, called PGSE-A and PGSE-B (consensus sequences are CCGGGGCGGGGCG and TTTTACAATTGGTC, respectively). However, transcription factors that bind to PGSEs have not yet been identified. Therefore, we attempted to identify these transcription factors in order to clarify the molecular mechanisms of the proteoglycan pathway of the mammalian Golgi stress response.

## Materials and Methods

### Cell culture and transfection

HeLa cells were cultured in Dulbecco’s modified Eagle’s medium (glucose at 4.5 g/liter) supplemented with 10% fetal calf serum, 2 mM glutamine, 100 units/ml of penicillin, and 100 g/ml of streptomycin at 37°C in a humidified 5% CO_2_, 95% air atmosphere (Yoshida *et al*., 1998). Cells were transfected with plasmid DNA by the calcium phosphate or lipofection method using FuGene 6 (Promega, WI) (Yoshida *et al*., 2000). After transfection, cells were treated with 7.5 mM 4MU-xylodside for 16-18 h, washed three times with PBS, and then harvested for immunoblotting, immunocytochemistry, and luciferase assays (Yoshida *et al*., 2001).

### Construction of plasmids and transfection of siRNAs

A 4xPGSE-A reporter plasmid was constructed by fusing the 4xPGSE-A sequence from the human NDST2 gene with firefly luciferase in the pGL4 vector (Sasaki et al., 2019). The expression plasmid for human SDC2 was kindly provided by Drs. Hiroshi Kitagawa and Satomi Nadanaka (Nadanaka and Kitagawa, manuscript in preparation). The plasmids expressing human KLF family proteins were purchased from Genscript (Piscataway, NJ) or OriGene (Rockville, MD). To construct promoter-luciferase reporter vectors, the corresponding promoter regions of the human KLF2 and KLF4 genes were amplified using human genomic DNA as a template and cloned into the BglII site of the GL4 basic vector (Promega, Fitchburg, WI). Point mutants of the vectors were constructed by site-directed mutagenesis using a QuikChange Site-Directed Mutagenesis kit (Stratagene, CA) (Yoshida *et al*., 2006).

### Immunoblotting and immunocytochemistry

Immunoblotting and immunocytochemistry were performed as previously reported (Uemura *et al*., 2009). Anti-myc, anti-β-tubulin, anti-GFP, anti-KLF2, and anti-KLF4 antisera were purchased from WAKO (#017-21871, Tokyo, Japan), WAKO (#014-25041, Tokyo, Japan), Santa Cruz Biotechnology (#sc-9996, Santa Cruz, TX), Abcam (#ab236507, Cambridge, UK), and Becton, Abcam (#ab215036), respectively.

### Quantitative RT-PCR (qRT-PCR) and luciferase assays

qRT-PCR was performed using a QuantStudio 6 Flex Real-Time PCR System (Thermo-Fischer Scientific, Waltham, MA) and PrimeScript RT reagent kit with gDNA Eraser and SYBR Premix Ex Taq II (Tli RNase H Plus) (TaKaRa, Kusatsu, Japan) (Taniguchi and Yoshida, 2017). The primer pairs used for qRT-PCR were as follows: *KLF2* (AAGACCTACACCAAGAGTTCGCATC and TCTGAGCGCGCAAACTTCC), *KLF4* (AAGAGTTCCCATCTCAAGGCACA and GGGCGAATTTCCATCCACAG), *KLF6* (CCACTTTAACGGCTGCAGGAA and CGGAAGTGCCTGGTTAACTCATC), *KLF7* (TTGTCCACGACACCGGCTAC and GGAAACAGTCCAAGTCCTCACCA), *NDST2* (CCTGTGA TGACAAGAGGCACAAA and TGCAGGCTCAGGAAGAAGTGAAT), *HS6ST1* (TCACCTTCACCA TGGGCTTC and GACTGAGACAAGACCCGTGCTTC), *GLCE* (GGCTACAATGTGGAAGTCCGAGA and CTGGATTGGATAGAAATAGCCTTGA), and *B3GAT3* (GAGCAGTCTTCTGAGCCACCTTG and CTGCTTCATCTTGGGCTTCTCTG). Assays for firefly luciferase were conducted with the PicaGene Dual Sea Pansy luminescence kit (Toyo Ink, Tokyo, Japan) (Uemura *et al*., 2013).

### Construction of KLF2/4-DKO cells

To establish KLF2/4-DKO cell lines, the KLF2 and KLF4 genes were knocked out in HeLa cells by the CRISPR-Cas9 method with sgRNAs (Fw 5’-caccgGTCGTCGGTGCCGCCGGACT-3’ and Rv 5’-aaacAGTCCGGCGGCACCGACGACc-3’ for KLF2; Fw 5’-caccgGTGGTGGCGCCCTACAACGG-3’ and Rv 5’-aaacCCGTTGTAGGGCGCCACCACc-3’ for KLF4). Genomic DNA was prepared from each KLF2/4-DKO cell line, and subjected to PCR using specific primers (5’-GTCCTTCTCCACTTTCGCCA-3’ and 5’-CAGCAGCTCAGACACCAGG-3’ for KLF2; 5’-GGTGAAGAAGGTGGGGTGAG-3’ and 5’-TTCAACCTGGCGGACATCAA-3’ for KLF4) in order to verify the disruption of the KLF2 and KLF4 genes. The disruption of each gene was confirmed by immunoblotting with anti-KLF2 and anti-KLF4 antisera.

## Results

### Screening of transcription factors that bind to PGSE-A *in silico*

We previously identified two enhancer elements, PGSE-A and PGSE-B, which were responsible for the transcriptional induction of the glycosylation enzymes of the proteoglycan pathway (Sasaki et al., 2019). To isolate transcription factors that bind to PGSE-A, we searched the JASPAR database (Castro-Mondragon *et al*., 2022) for transcription factors with similar predicted binding sites to the nucleotide sequence of PGSE-A (CCGGGGCGGGGCG), and found that the binding sites of KLF family transcription factors (KLFs) resembled the PGSE-A sequence (Fig. 1). KLFs consisted of 17 proteins (KLF1 – KLF17), and 14 KLF proteins were included in the top 24 transcription factors in this *in silico* screening. These results suggest the potential of KLFs as the transcription factors responsible for transcriptional induction from PGSE-A.

**Fig. 1.**
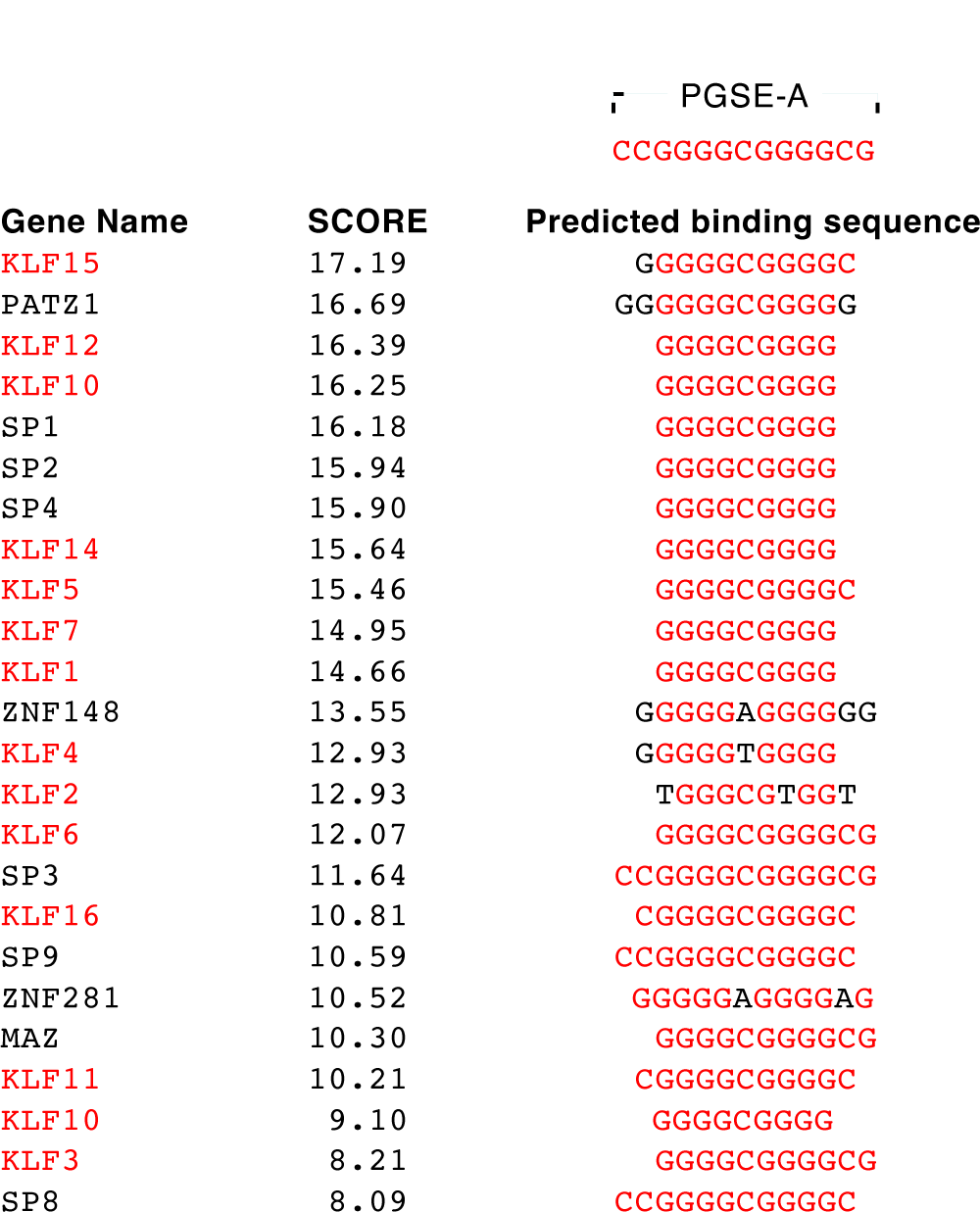
*In silico* screening of PGSE-A-binding proteins using the JASPAR database. The top 24 transcription factors of which the predicted binding sequences are similar to the PGSE-A sequence are shown. SCORE is a value calculated by JASPAR. Nucleotides identical to the PGSE-A consensus sequence are shown in red.

### Overexpression of KLF2 or KLF4 enhances transcription from PGSE-A

We then investigated whether the overexpression of KLFs affected transcription from PGSE-A (Fig. 2). HeLa cells transfected with an expression vector of KLFs as well as a PGSE-A reporter plasmid were treated with 4MU-xyloside (a PG-type Golgi stress inducer) and subjected to immunoblotting to confirm the expression of KLFs (Fig. 2A and 2B) and a luciferase assay to evaluate the effects of the overexpression of KLFs (Fig. 2C and 2D). KLF2 and KLF4 were the most effective among KLFs (Fig. 2C, lanes 13 and 15). The amino acid sequences of the DNA-binding motifs of KLF2 and KLF4 were quite similar among KLFs (only seven amino acid residues were replaced) (Fig. 3), suggesting that KLF2 and KLF4 bind to similar nucleotide sequences, such as PGSE-A.

**Fig. 2.**
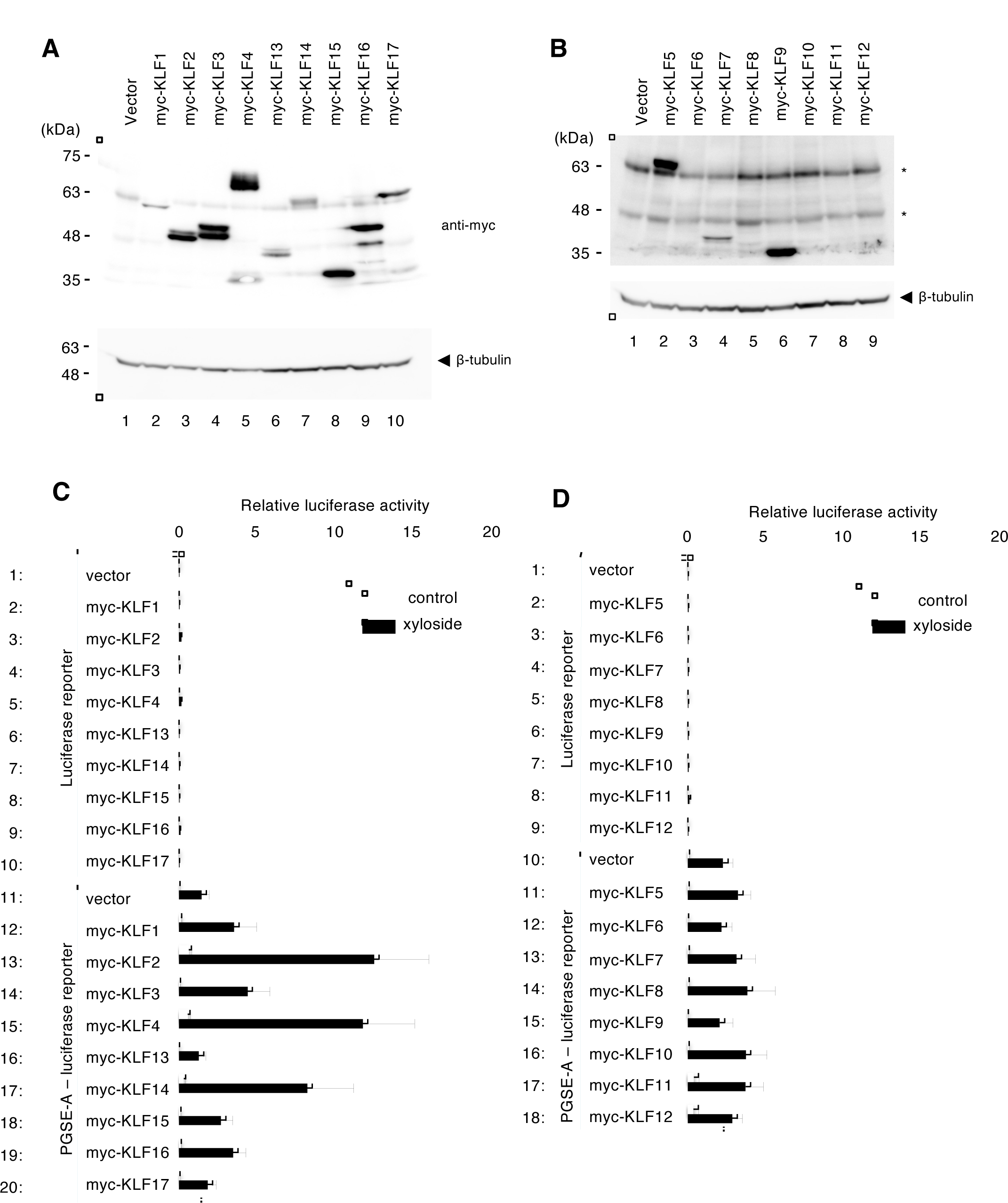
Effects of the overexpression of KLF family transcription factors on transcriptional induction from PGSE-A. Expression check of myc-KLF (A and B) and effects of myc-KLFs overexpression on a PGSE-A reporter (C and D). HeLa cells were co-transfected with an expression vector of myc-KLFs and a PGSE-A-Luciferase reporter, and subjected to immunoblotting (A and B) and a luciferase assay (C and D). Asterisks indicate non-specific bands. Experiments in (C) and (D) were performed in triplicate.

**Fig. 3.**
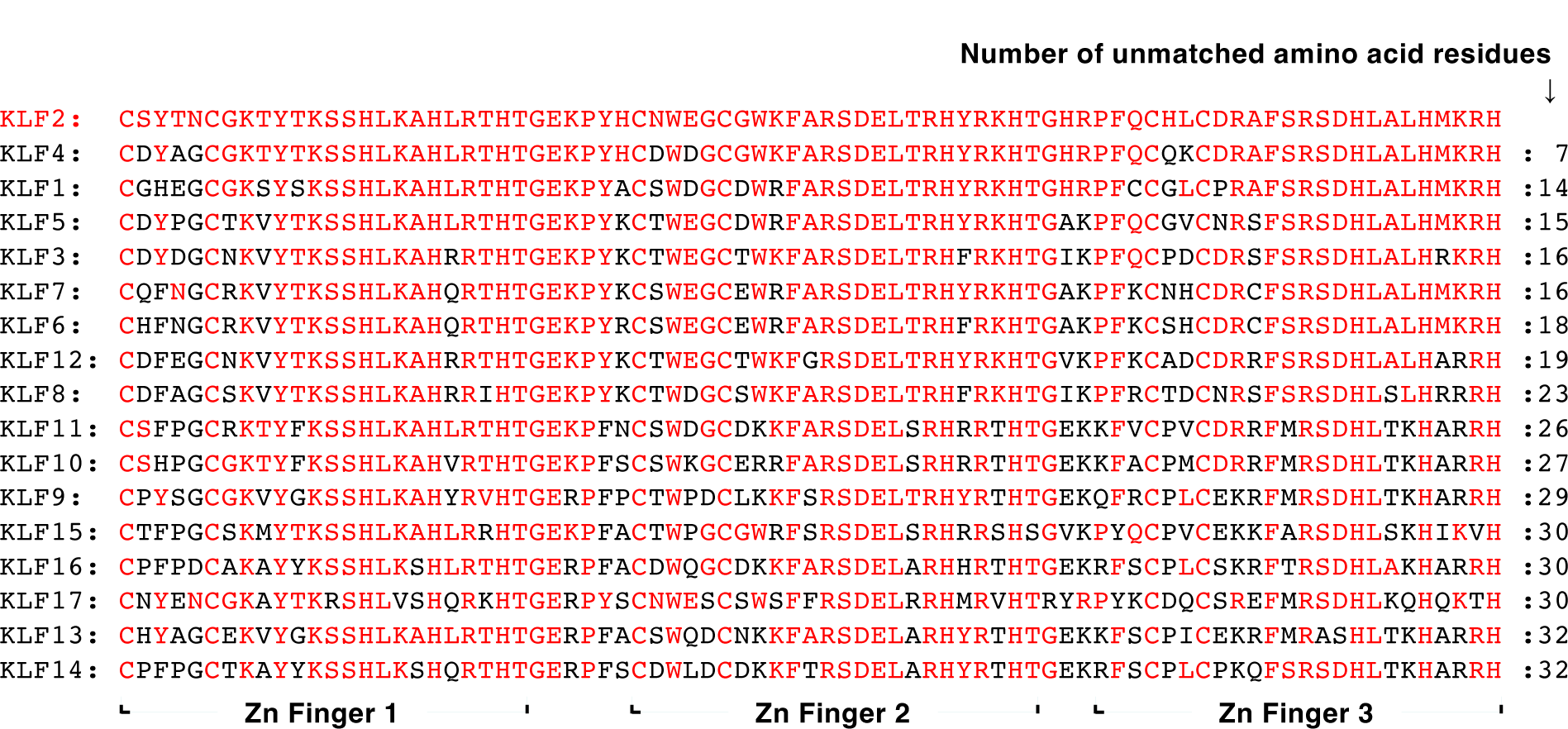
Comparison of DNA-binding domains of human KLF proteins. Amino acid residues identical to those of the KLF2 protein are shown in red.

### Expression of dominant negative (DN) mutants of KLF2 or KLF4 suppresses transcriptional induction from PGSE-A upon PG-type Golgi stress

We investigated whether the expression of DN mutants of KLF2 and KLF4 suppressed transcriptional induction from PGSE-A (Fig. 4). We produced two types of DN mutants that lacked the activation domain (KLF2 (108-355) and KLF4 (171-513)) or the activation and inhibitory domains (KLF2 (251-355) and KLF4 (400-513)) (Fig. 4A and 4B). HeLa cells transfected with an expression plasmid of DN mutants and a PGSE-A reporter were treated with 4MU-xyloside, and the luciferase activity of cell lysates was measured (Fig. 4C and 4D). We found that the DN mutants of KLF2 and KLF4 both significantly reduced increases in luciferase activity upon the 4MU-xyloside treatment, indicating that these DN mutants suppress transcriptional induction from PGSE-A, possibly by preventing endogenous transcription factors from binding to PGSE-A. We examined the expression of these DN mutants by immunoblotting (Fig. 4E and 4F). We also tried to investigate whether transcriptional induction from PGSE-A was abolished in KLF2 and KLF4 double knockout (KLF2/4-DKO) cells, but we were not able to establish KLF2/4-DKO cell lines using the CRISPR-Cas9 system. It may be difficult to disrupt both alleles of the KLF2 and KLF4 genes since they are important for cell growth.

**Fig. 4.**
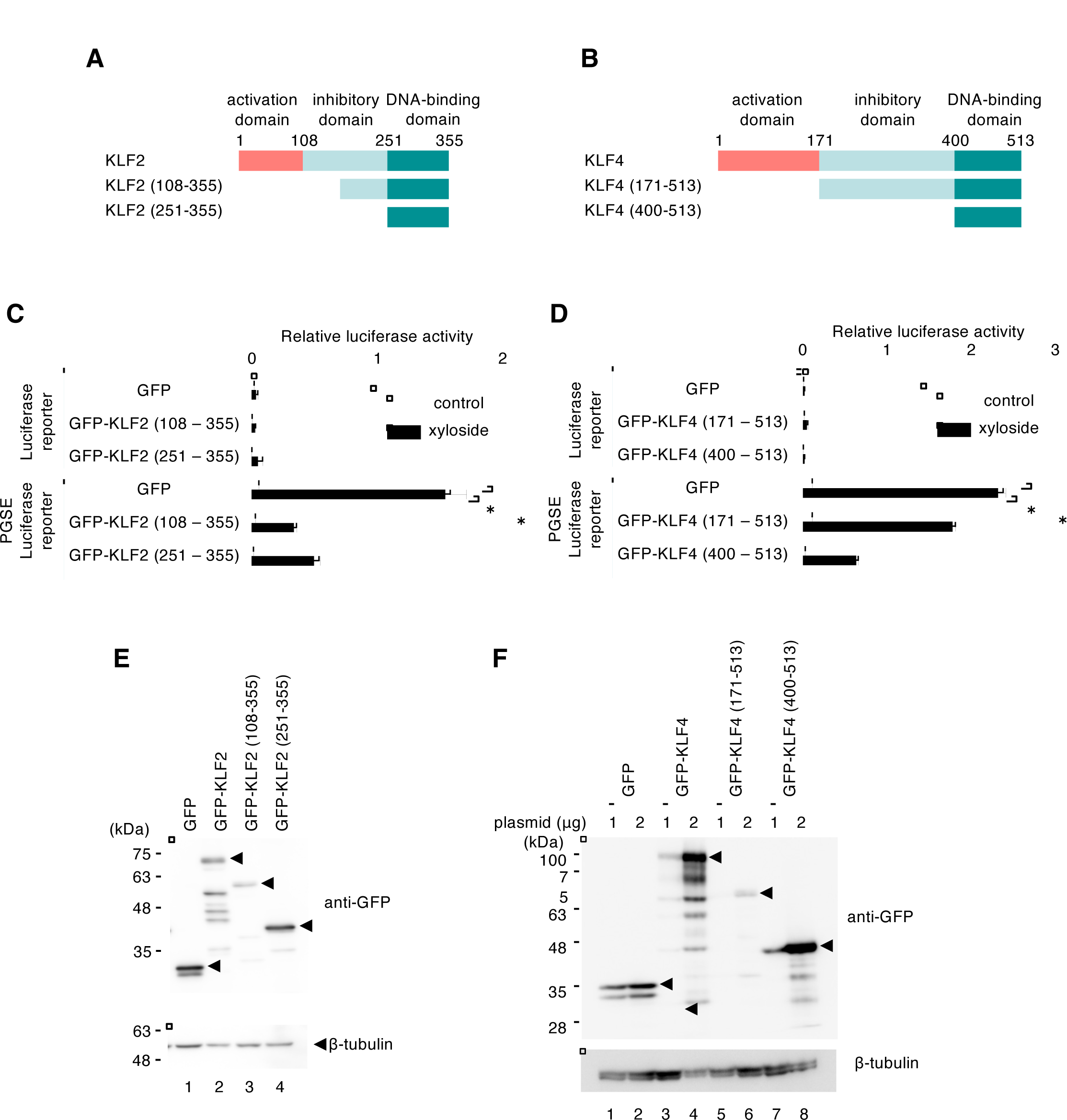
Effects of dominant negative mutants of KLF2 and KLF4 on transcriptional induction from PGSE-A. (A and B) Structures of dominant negative mutants of KLF2 and KLF4. (C and D) The effects of KLF2 and KLF4 dominant negative mutants on a PGSE-A reporter were analyzed as described in Fig. 2. Values are the means ± SE of three independent experiments. ***, P < 0.001; **, P < 0.01; *, P < 0.05. (E and F) Expression of KLF2 and KLF4 dominant negative mutants. (E and F) Expression check of each dominant negative mutant by immunoblotting.

### Transcription of KLF2 and KLF4 genes is induced upon PG-type Golgi stress

Next we examined whether the expression of the KLF2 and KLF4 proteins was changed during PG-type Golgi stress (Fig. 5). HeLa cells were treated with 4MU-xyloside and MG132 (a proteasome inhibitor) and subjected to immunoblotting. The expression of KLF2 and KLF4 was increased by the treatment with 4MU-xyloside and MG132 in wild type cells (lanes 3 and 4), suggesting that the expression of these transcription factors was induced by PG-type Golgi stress and that they were rapidly degraded by the ubiquitin-proteasome system.

**Fig. 5.**
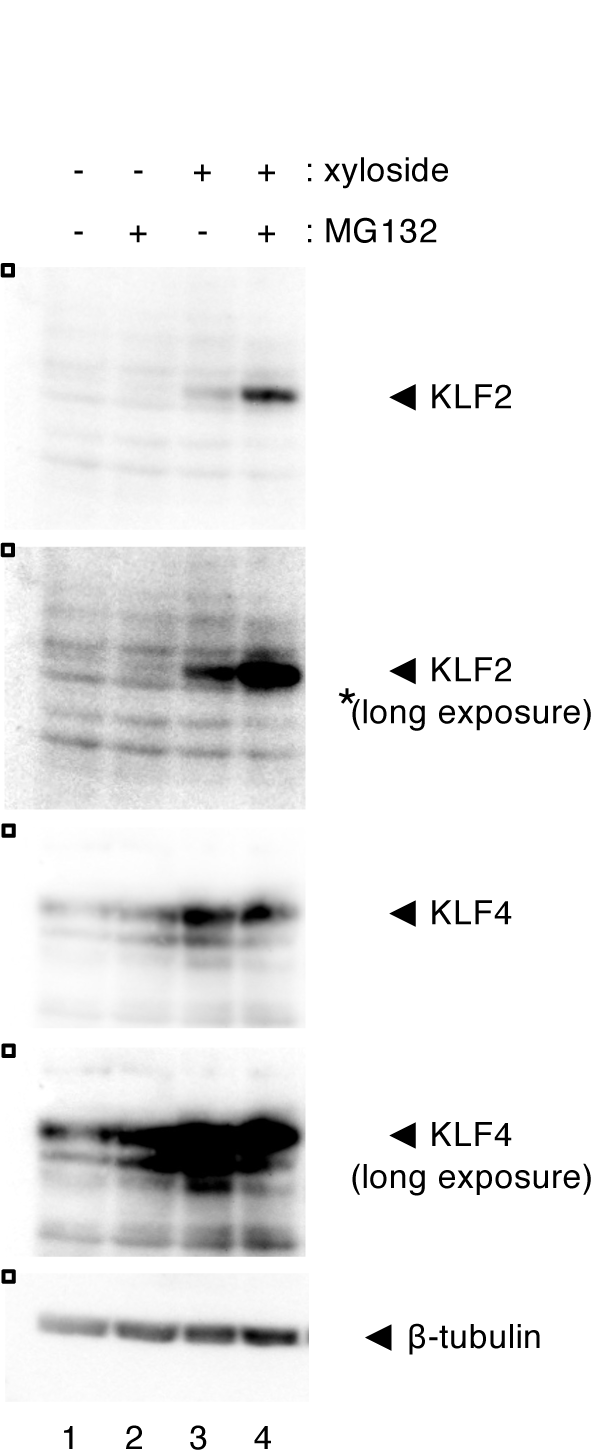
Expression of KLF2 and KLF4 proteins during PG-type Golgi stress. HeLa cells treated with 7.5 mM 4MU-xyloside for 18 h and with 10 μM MG132 for 2 h were analyzed by immunoblotting using anti-KLF2 and anti-KLF4 antisera.

Since the expression of the KLF2 and KLF4 proteins was increased upon PG-type Golgi stress (Fig. 5), we investigated whether the expression of their mRNAs was up-regulated in response to this stress. Data from previous RNA sequencing analyses (Sasaki et al., 2019) indicated that the mRNA levels of KLF2, KLF4, KLF6 and KLF7 were significantly increased by a treatment with 4MU-xyloside (Fig. 6A, lanes 2, 4, 6 and 8). We also examined their mRNA levels by qRT-PCR. Total RNA was prepared from HeLa cells treated with 4MU-xyloside and subjected to qRT-PCR analyses (Fig. 6B). The expression of these mRNA was increased by the treatment with 4MU-xyloside (lanes 2, 4, 6 and 8). Time-course experiments revealed that the induction pattern of KLF2 and KLF4 mRNAs was similar to that of glycosylation enzymes such as HS6ST1 and NDST2 (Fig. 6C).

**Fig. 6.**
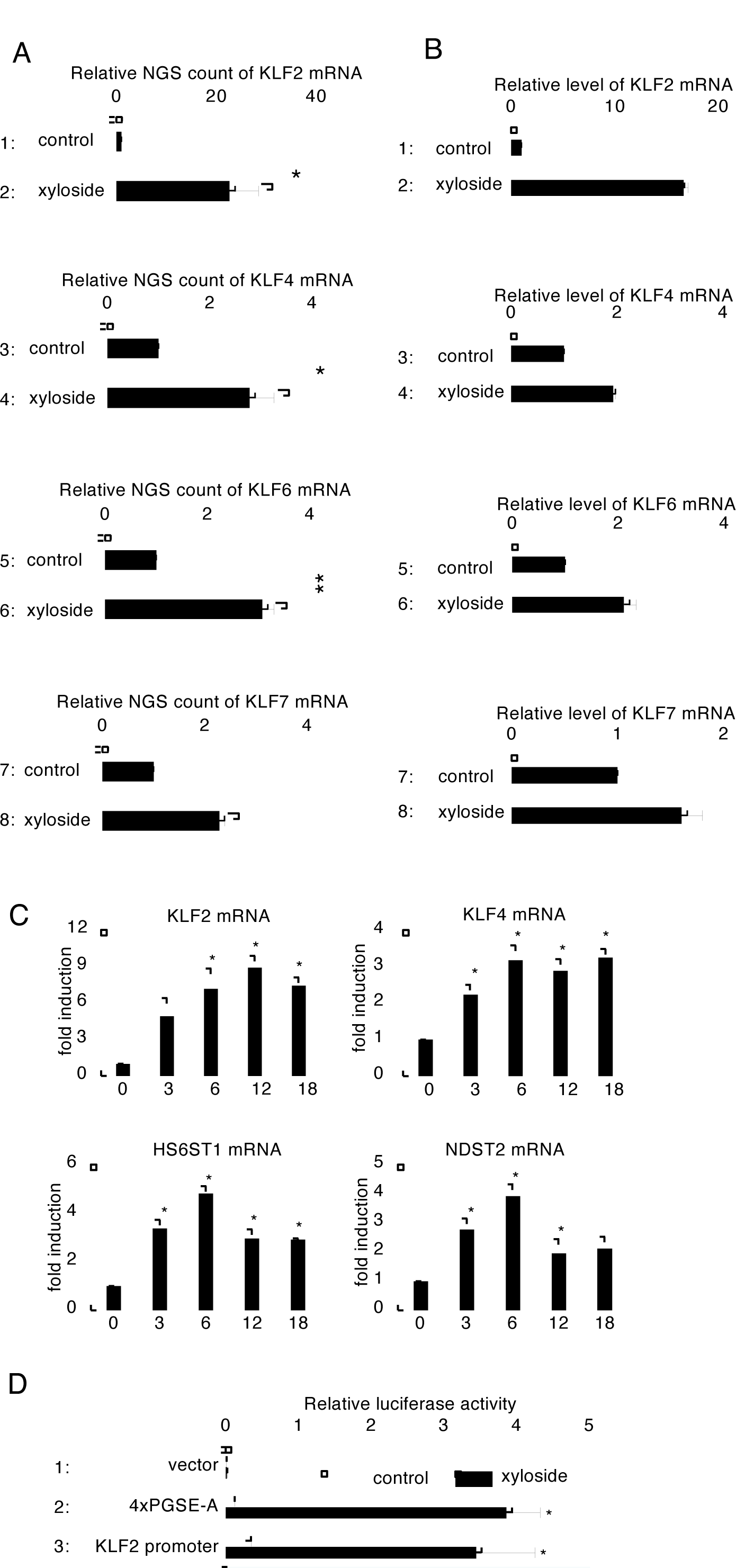
Expression of KLF2 and KLF4 mRNAs during PG-type Golgi stress. (A) RNA sequencing analysis of KLF family genes. Total RNA prepared from HeLa cells treated with 7.5 mM 4MU-xyloside for 18 h was subjected to RNA sequencing. (B) Results of a qRT-PCR analysis of KLF family genes. Total RNA prepared in (A) was subjected to a qRT-PCR analysis. (C) Induction time-course of the mRNAs of KLF2, KLF4, and glycosylation enzymes. Total RNA prepared from HeLa cells treated with 7.5 mM 4MU xyloside for the indicated time was subjected to qRT-PCR. (D) Luciferase analysis of the human KLF2 promoter. The [−961 to +140] region of the human KLF2 promoter was fused with the firefly luciferase reporter gene and subjected to a luciferase assay with 7.5 mM 4MU-xyloside. Experiments were repeated four times. Reporter plasmid pGL4 basic and 4xPGSE-A reporter were used as the negative and positive controls, respectively.

We then investigated whether transcription from the promoters of human KLF2 and KLF4 was induced upon PG-type Golgi stress (Fig. 6D and 9A). The [−961 to +140] and [−932 to +132] regions of the human KLF2 and KLF4 promoters were fused with firefly luciferase and transfected into HeLa cells. When cells were treated with 4MU-xyloside, the expression of luciferase significantly increased (Fig. 6D, lane 3 and Fig. 9A, lane 2), indicating that the transcription of the human KLF2 and KLF4 genes was induced by PG-type Golgi stress.

### Identification of enhancer elements responsible for the transcriptional induction of the KLF2 gene

We attempted to identify the enhancer elements regulating transcriptional induction from the KLF2 promoter in response to PG-type Golgi stress. The [−961 to +140] region of the human KLF2 promoter contained two PGSE-A-like sequences (PGSE-A1 and PGSE-A2 in Fig. 7A), suggesting that these PGSE-A-like sequences were responsible for the transcriptional induction of the KLF2 gene in response to Golgi stress. When PGSE-A1 and PGSE-A2 in the human KLF2 promoter were mutated, transcriptional induction from the KLF2 promoter only slightly decreased (Fig. 7F, lanes 3 and 4), indicating that these PGSE-A-like sequences were not the main regulators. To identify other enhancers regulating the transcriptional induction of the KLF2 gene, we generated serial deletion mutants of the KLF2 promoter and evaluated transcriptional induction from these deletion mutants, similar to that shown in Fig. 5E (Fig. 7B). The results obtained revealed that the [−135 to +140] region of the KLF2 promoter showed notable transcriptional induction (lane 5), whereas the [−99 to +140] region exhibited very small induction (lane 6), suggesting that the [−135 to −99] region contained strong enhancers. We produced another set of deletion mutants, evaluated their activities, and found that the [−135 to −119] region was important for transcriptional induction (Fig. 7C, lanes 2 and 3). Point mutations were introduced into this region, and the TTATATA sequence was found to be crucial for transcriptional induction (Fig. 7D, lanes 3-9).

**Fig. 7.**
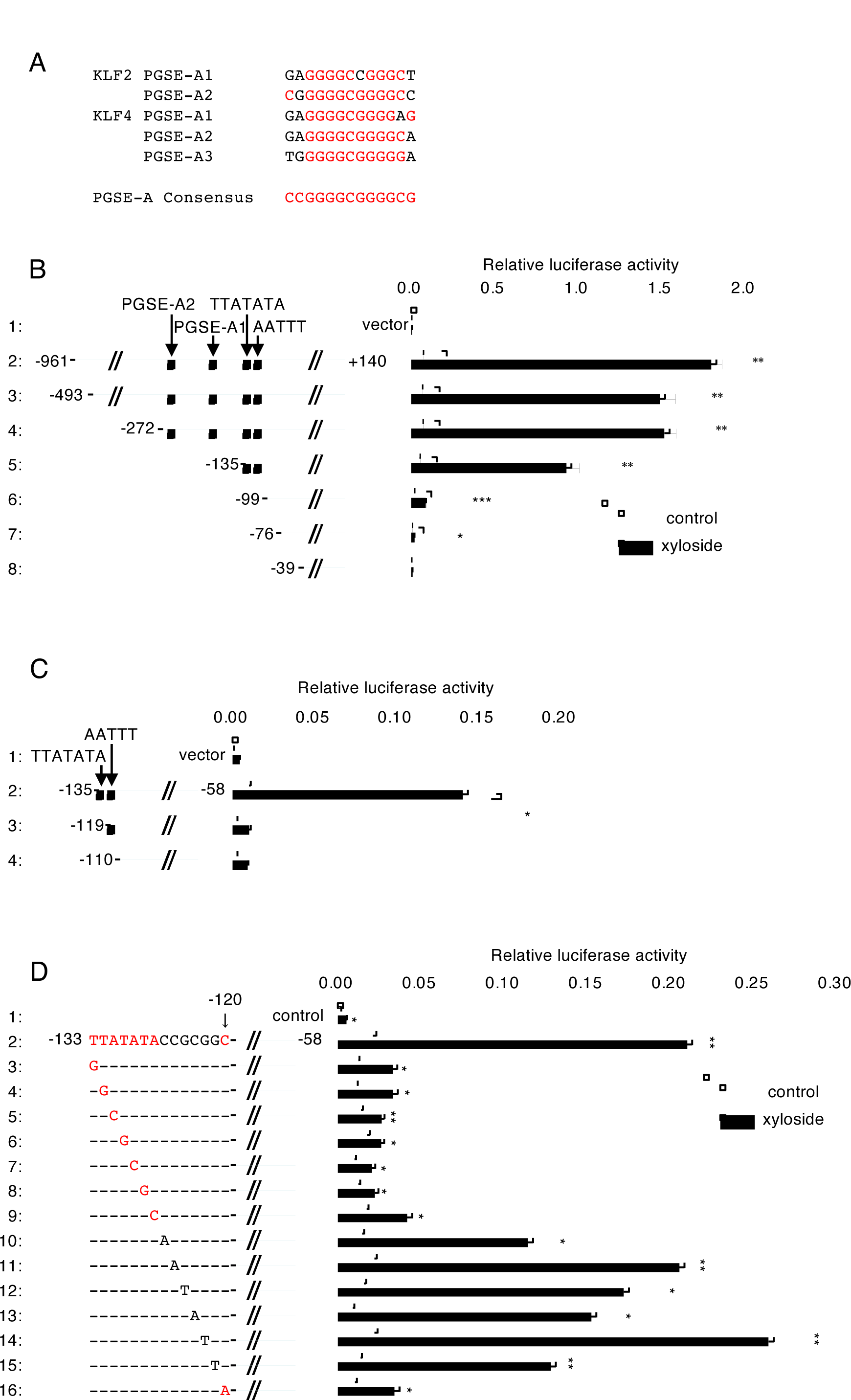
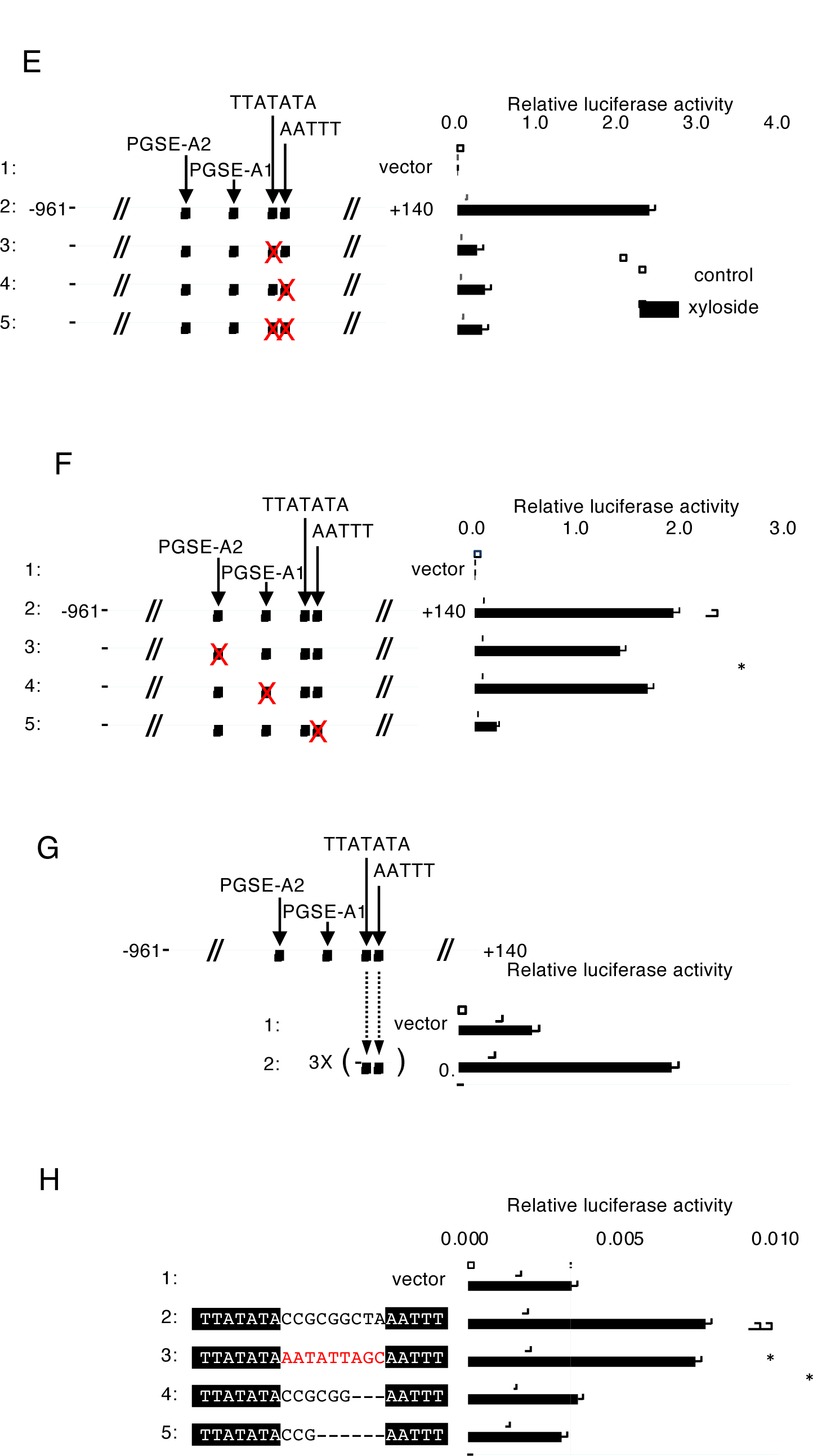
Analysis of the human KLF2 promoter. (A) Comparison of PGSE-A like sequences in the human KLF2 and KLF4 genes. (B and C) Effects of 5’ and 3’ deletions of the KLF2 promoter on transcriptional induction. The indicated deletion mutants of the KLF2 promoter fused with the luciferase reporter were subjected to a luciferase assay, as shown in Fig. 6D. Experiments were performed in triplicate. (D) Point mutation analysis of the [−133 to −58] region of the KLF2 promoter. Each nucleotide in the region was replaced with another nucleotide in transversion, and transcription induction from the resultant point mutants was evaluated by a luciferase assay, as described in (B). Mutations in nucleotides that markedly reduced transcriptional induction were indicated in red. Experiments were performed in triplicate. (E and F) Effects of mutations in the TTATATA, AATTT, and PGSE-A like sequences on transcriptional induction. Transcriptional induction from the [−961 to +140] region with the indicated mutations was evaluated as described in (B). (G) Transcriptional induction from the TTATATA(N9)AATTT sequences. Three repeats of the TTATATA(N9)AATTT sequences were fused with the luciferase reporter, and their transcriptional induction activity was evaluated, as described in (B). Experiments were performed in triplicate. (H) Stepwise deletion mutation in the spacer sequence between the TTATATA and AATTT sequences. Cells transfected with the indicated reporter plasmids were processed as described in (B). Reporter plasmid pGL4 basic was used as the control. Experiments were performed in triplicate.

Jain and colleagues previously reported that the [−135 to −99] region contained a single consensus MEF-binding site (AATTT) that was typically bound by members of the MADS-box family of transcriptional regulators (Sen-Banerjee *et al*., 2005). When this AATTT sequence was mutated in the KLF2 promoter, transcriptional induction markedly decreased (Fig. 7E, lane 4), suggesting that the AATTT sequence is an important enhancer of the transcriptional induction of KLF2 in response to Golgi stress. Mutations in the TTATATA sequence also showed reductions in transcriptional induction similar to that of the AATTT sequence (lane 3). The simultaneous mutation of these two sequences did not result in additive effects (lane 5), indicating that these sequences worked together. We then evaluated the activities of the TTATATA and AATTT sequences alone (Fig. 7G). Three copies of the [−135 to −111] region containing the TTATATA and AATTT sequences showed strong transcriptional induction (lane 2), which suggested that the TTATATA sequence and AATTT sequence was sufficient for transcriptional induction. We also examined the importance of the spacer region between the TTATATA sequence and AATTT sequence (Fig. 7H). When all nucleotides were mutated in the spacer region (the [−135 to −111] region), transcriptional induction was not affected (lane 3), indicating that the nucleotide sequence in the spacer region was not important. When the distance between the TTATATA and AATTT sequences was changed, transcriptional induction was significantly reduced (lanes 4 and 5), suggesting that the distance between the TTATATA and AATTT sequences was critical. These results indicate that the TTATATA sequence cooperated with the AATTT sequence to enhance the transcription of the KLF2 gene. Therefore, we named the TTATATA(N9)AATTT sequence PGSE-C. Comparisons of PGSE-C and the surrounding sequences in the KLF2 promoters of various vertebrates revealed that the PGSE-C sequence was well conserved during evolution (Fig. 8A).

**Fig. 8.**
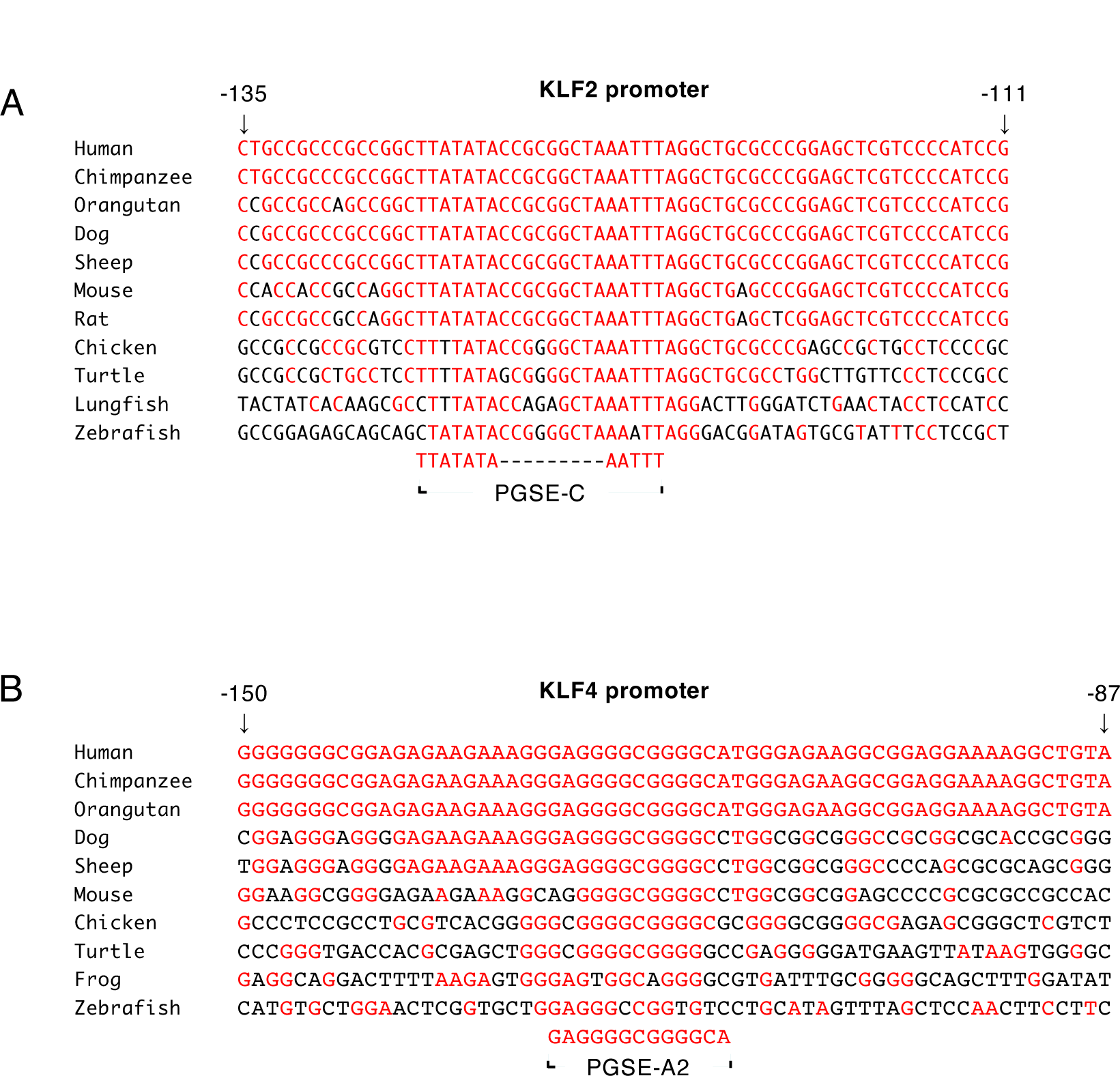
Conservation of PGSE-C in the vertebrate KLF2 promoter and PGSE-A2 in the KLF4 promoter during evolution. PGSE-C sequences in the human KLF2 promoter (A) as well as PGSE-A2 sequences in the human KLF4 promoters in various vertebrate animals (B) were aligned. Conserved nucleotides are indicated in red. Animals are *Pan troglodytes* (Chimpanzee), Pongo *Lacépède* (Orangutan), *Canis lupus familiaris Ovis aries* (Sheep), *Dermochelys coriacea* (leatherback sea turtle), *Xenopus tropicalis* (tropical clawed frog), and *Danio rerio* (zebrafish).

### Identification of enhancer elements responsible for the transcriptional induction of the KLF4 gene

We attempted to identify the enhancers regulating the transcriptional induction of the KLF4 gene (Fig. 9). The [−932 to +132] region of the human KLF4 gene and its deletion mutants were fused with the luciferase reporter gene and transfected into HeLa cells in order to evaluate its transcriptional induction activity upon a xyloside treatment (Fig. 9A). The luciferase assay showed that the [−932 to +132] and [−367 to +132] regions both exhibited strong activity (lanes 2 and 3), while further deletions significantly reduced this activity (lanes 4-6), suggesting that strong enhancers of Golgi stress-induced transcription resided in the [−367 to +132] region. We found that the [−367 to +132] region contained three PGSE-A-like sequences (PGSE-A1, PGSE-A2 and PGSE-A3) (Fig. 7A). When these PGSE-A-like sequences were mutated, the KLF4 promoter almost lost its transcriptional induction activity (Fig. 9B, lanes 3-5). These results indicate that the transcriptional induction of the KLF4 gene was mainly regulated by PGSE-As, particularly PGSE-A2. Comparisons of PGSE-A2 and the surrounding sequences in the KLF4 promoters of various vertebrates suggested that the PGSE-A2 sequence was well conserved during evolution (Fig. 8B), indicating the importance of PGSE-A2.

**Fig. 9.**
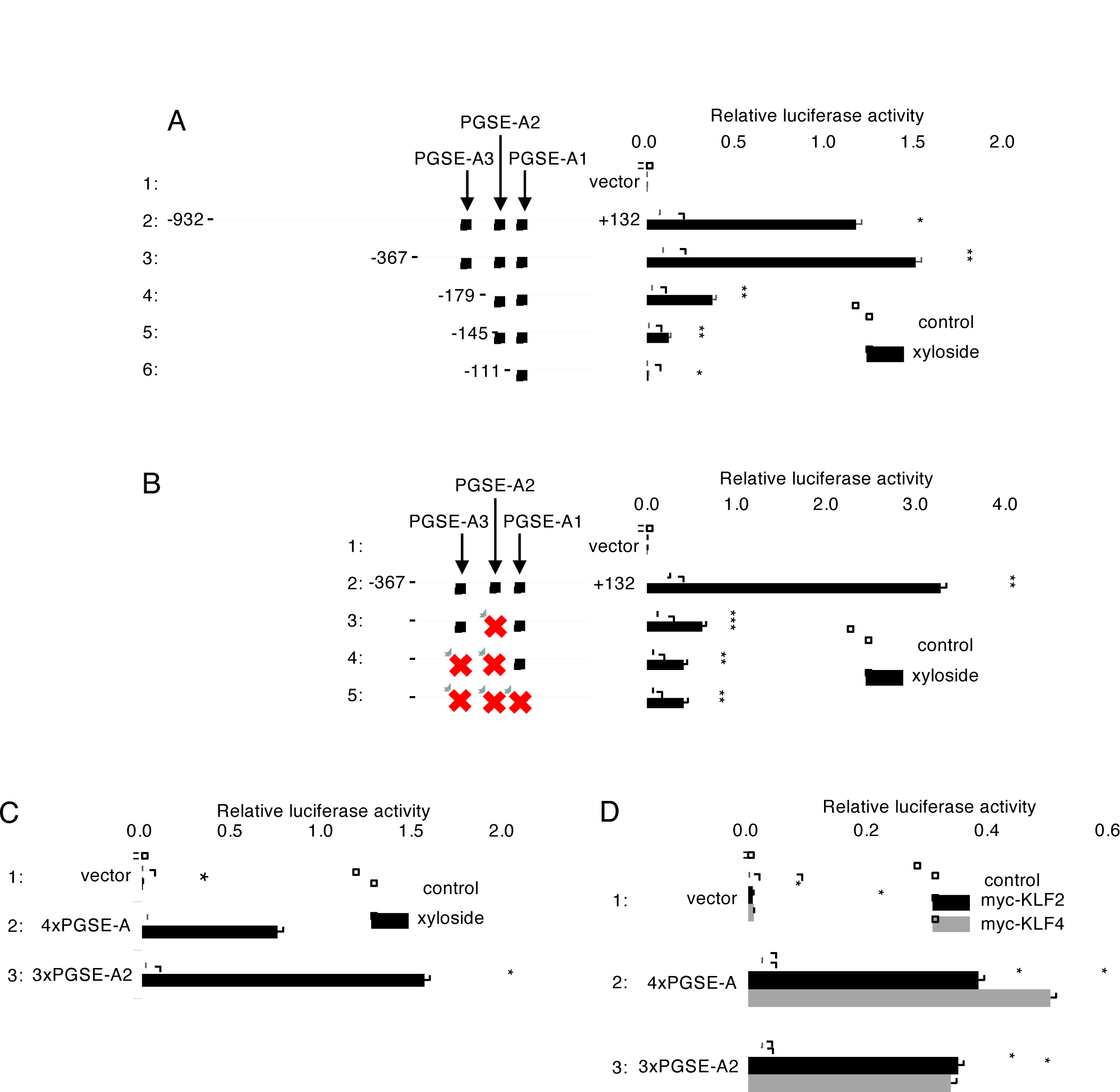
Analysis of the human KLF4 promoter. (A) Transcriptional induction from 5’ deletion mutants of the human KLF4 promoter. The indicated deletion mutants were fused with the luciferase gene and their transcriptional induction activity was evaluated as described in Fig. 6D. Experiments were performed in triplicate. (B) Transcriptional induction from PGSE-mutants of the KLF4 promoter. The indicated PGSE-mutants were subjected to a luciferase assay as described in (A). Experiments were repeated four times. (C and D) Transcriptional induction activity of PGSE-A2 in the KLF4 promoter upon the 4MU-xyloside treatment (C) and KLF2 and KLF4 overexpression (D). Three repeats of the PGSE-A2 sequence fused with the luciferase gene were analyzed by a luciferase assay as described in (A). Experiments were performed in triplicate. A 4xPGSE-A reporter was used as a positive control.

## Discussion

We herein identified KLF2 and KLF4 as candidate transcription factors that bind to the PGSE-A enhancer and regulate transcriptional induction by the proteoglycan pathway of the mammalian Golgi stress response. The expression of KLF2 and KLF4 was up-regulated in response to PG-type Golgi stress. Moreover, we identified the enhancer elements regulating the transcriptional induction of the human KLF2 and KLF4 genes, namely, PGSE-C and PGSE-A, respectively. Based on the present results, we propose the working hypothesis shown in Fig. 10. (1) Upon PG-type Golgi stress, unidentified transcription factors are activated, bind to PGSE-C, and induce the transcription of the KLF2 gene, leading to the up-regulated expression of KLF2. (2) KLF2 binds to the PGSE-A sequence in the promoters of the KLF2, KLF4 and glycosylation enzymes for proteoglycans. (3) KLF4 induced by KLF2 further enhances transcription from PGSE-A, (4) while induced glycosylation enzymes relieve Golgi stress.

**Fig. 10.**
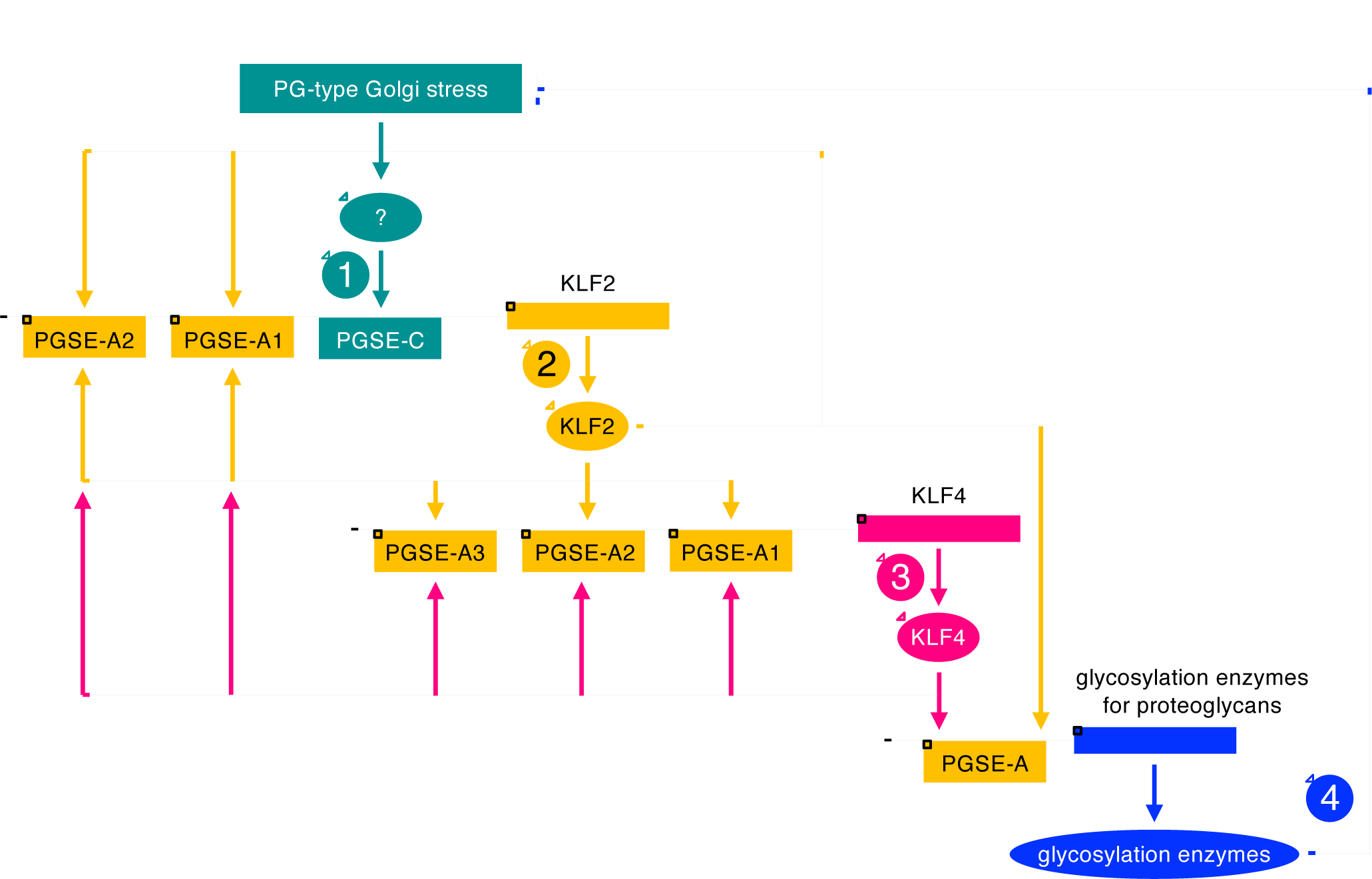
Working hypothesis of the proteoglycan pathway of the mammalian Golgi stress response. Upon PG-type Golgi stress, an unknown transcription factor induces the transcription of the KLF2 gene through PGSE-C (indicated in green), leading to an increase in the expression of KLF2. KLF2 increases the transcription of KLF2, KLF4, and glycosylation enzymes for proteoglycans (indicated in yellow). KLF4 enhances the transcription of the same set of genes (indicated in red). The induced glycosylation enzymes then relieve Golgi stress (indicated in blue).

KLF2 and KLF4 belong to the Krüppel-like family of transcription factors (KLFs), which contain three conserved C2H2 zinc finger DNA-binding domains (Oishi and Manabe, 2018). There are 17 KLF genes in the human genome: KLF1-17. KLF2/LKLF (lung KLF) was initially isolated as a transcription factor similar to KLF1/EKLF (erythroid KLF), and is predominantly expressed in the lungs (Anderson *et al*., 1995). Knockout mice of KLF2 died between E12.5 and E14.5 from severe intra-embryonic and intra-amniotic hemorrhaging due to a defect in the assembly of the vascular tunica media and concomitant vessel wall stabilization during mammalian embryogenesis, partly because of a decrease in the deposition of the extracellular matrix in blood vessels (Kuo *et al*., 1997). The induced expression of glycosylation enzymes for proteoglycans may be less in the blood vessels of KLF2 knockout mice, which decreased the deposition of proteoglycans and vessel wall destabilization. KLF4/GKLF (gut KLF) was initially identified as a KLF family protein, the expression of which is enriched in the gut (Shields *et al*., 1996). KLF4 is one of the Yamanaka factors that are required for the induction of pluripotent stem cells (Takahashi and Yamanaka, 2006). The DNA-binding motifs of KLF2 are similar to those of KLF4 (Fig. 1B), and KLF2 and KLF4 have been shown to jointly control endothelial identity and vascular integrity (Sangwung *et al*., 2017), suggesting that the functions of KLF2 and KLF4 are redundant. KLF2 and KLF4 may enhance the formation of endothelial glycocalyx by up-regulating the expression of the proteoglycan-type of glycosylation enzymes in response to Golgi stress in endothelial cells, and maintain the strength of blood vessels. We herein showed that the overexpression of KLF2 or KLF4 enhanced transcriptional induction from PGSE-A (Fig. 2), while that of their DN mutants suppressed transcriptional induction from PGSE-A (Fig. 3), which is consistent with this hypothesis. KLF proteins other than KLF2 and KLF4 may be involved in residual transcriptional induction from PGSE-A. The present results showing that the expression of KLF6 and KLF7 mRNA was induced by PG-type Golgi stress (Fig. 6A and 6B) coincides with this notion.

We identified a novel enhancer element PGSE-C with the consensus sequence TTATATA(N9)AATTT, which is responsible for the transcriptional induction of the human KLF2 gene upon PG-type Golgi stress. This enhancer is well conserved during evolution (Fig. 7A), suggesting the importance of PGSE-C for the transcriptional induction of KLF2 genes. We also found that PGSE-As were responsible for the transcriptional induction of the human KLF4 gene, to which PGSE-A2 is the main contributor. The sequence of PGSE-A2 was also highly conserved in vertebrates (Fig. 7B), suggesting the importance of PGSE-A2 for the transcriptional induction of the KLF4 genes in vertebrates. We searched for transcription factors binding to PGSE-C using the JASPAR database, and obtained several candidates, including FOXO family transcription factors (manuscript in preparation).

In conclusion, we herein revealed an important aspect of the complex mechanism of the mammalian Golgi stress response, which is one of the fundamental issues in cell biology.

## Acknowledgments

We thank the members of JSPS Grant “Organelle Zones” for their critical discussions, Dr. Hiroshi Kitagawa and Dr. Satomi Nadanaka for providing the SDC2 expression vector (pCMV-SDC2-FLAG), and Ms. Mikiko Ochiai and Ms. Miyu Sakamoto for their secretarial and technical assistance. This work was supported by JSPS KAKENHI (Grant numbers JP17K15122, JP17J00067, and JP22K06208, a Grant-in-Aid for Scientific Research on Innovative Areas of MEXT (JP17H06414) “Organelle Zones”, The Takeda Science Foundation, The Foundation of Kinoshita Memorial Enterprise, AMED-CREST “Proteostasis” and Nanken-Kyoten (Grant No. 2022-Domestic 08), TMDU.

## Notes

### Competing Interest Statement

The authors have declared no competing interest.

